# Myonuclear maturation dynamics in aged and adult regenerating mouse skeletal muscle

**DOI:** 10.1101/2021.07.13.452218

**Authors:** Jesse V. Kurland, Ashleigh Van Deusen, Brad Pawlikowski, Monica Hall, Nicole Dalla Betta, Tiffany Antwine, Alicia C Cutler, Alan Russell, Mary Ann Allen, Robin Dowell, Bradley Olwin

## Abstract

Skeletal muscle cells are multinucleated syncytial cells arising from cell fusion, yet despite sharing a common cytoplasm individual myonuclei express distinct transcriptional programs. Whether individual myonuclei acquire heterogenous transcriptional states via differences in their progenitors, during differentiation, or once their anatomical position is acquired, is not known. We performed transcriptome and pseudotime analysis of single myogenic nuclei from uninjured and post-injury murine skeletal muscle to assess when myonuclear heterogeneity is acquired. Two distinct progenitors contribute to myonuclei, one a non-myogenic fibroblast subtype, and skeletal muscle stem cells the other. Both progenitors enter a single pseudotime trajectory that bifurcates as myonuclei mature into two branches segregated by myosin isoform expression and metabolic profiles, suggesting transcriptional heterogeneity is acquired as myonuclei mature. In aged skeletal muscle myogenic progenitor expansion is perturbed and nuclei from aged muscle display distinct pseudotemporal kinetics compared to nuclei from young mice. In aged mice, the inferred myogenic differentiation trajectory is delayed, altering the distribution of myogenic nuclei in pseudotime, suggesting that altered transcriptional dynamics in nuclei in aged mice may drive age-associated muscle deficits and bias myonuclei towards acquiring oxidative metabolic profiles.

## Introduction

Skeletal muscle, critical for overall organismal health, deteriorates with age leading to reduced mobility and increased morbidity (1–3). Skeletal muscle myofibers are large multinucleated cells formed from fusion of skeletal muscle progenitors during development (4, 5). Within myofibers, mature myonuclei are transcriptionally distinct and express spatially defined sets of genes enabling the formation of key anatomical structures such as the neuromuscular junction (NMJ) or myotendinous junction (MTJ) (6–12). Although extensive transcriptional heterogeneity exists even beyond anatomically-defined genes, much is unknown regarding how heterogeneity in myonuclear gene expression is acquired (7, 8, 12–14).

Myonuclear transcribed genes include myosin heavy-chain (*Myh*) isoforms whose expression varies across muscle types, organisms’ age, and is heterogeneous even within individual myofibers (7, 15, 16). Myh proteins dictate sarcomere contractile speed, identifying myofiber type in mature muscle (17). During development and regeneration, new terminally differentiated immature myonuclei sequentially express embryonic *Myh3*, followed by neonatal *Myh8* as they mature, with adult myosin heavy chain isoforms replacing *Myh8* in mature myonuclei (16). *Myh7* expression defines slow twitch type-I muscle, while *Myh1* (type-IIx), *Myh2* (type-IIa), and *Myh4* (type-IIb) specify fast twitch type-II myofibers (17). A decrease in fast twitch myofibers is a hallmark of aged muscle, but the causes of this phenotype are incompletely understood (18).

Despite the presence of resident muscle stem cells (MuSCs) required to maintain and repair skeletal muscle, MuSCs lose their regenerative capacity during aging, likely contributing to reduced muscle function, reduced muscle size, and compromised innervation (19–26). We utilized regenerating muscle to probe temporal changes in myonuclear production in young adult and aged mice and derive a pseudotime trajectory of myonuclear differentiation. The ensuing trajectory imparts a continuous path of differentiation from myogenic progenitors into mature myonuclei, enabling the comparison of myonuclear maturation after an injury in young adult and aged murine skeletal muscle.

## Results

### EdU labeling of aged and adult mice reveals timing of MuSC expansion after an injury

We assessed the kinetics of MuSC expansion and decline following an injury to the TA muscles of young adult and aged mice to identify relevant timepoints for evaluating gene expression changes occurring during myonuclear maturation. MuSCs (Pax7+) exit quiescence and their myogenic progenitor progeny (myoblasts) expand rapidly, reaching a peak number by 4 days post-injury (dpi) in young adult mouse muscle, and slowly decline to within 2-fold of the initial MuSC numbers by 28 dpi (Fig. 1A). In aged mice expansion is delayed with myoblast numbers never reaching a comparable peak to that observed in young adult muscle (Fig. 1A). However, after 7 dpi the numbers of MuSCs and myoblasts in young adult and aged mice are indistinguishable, and by completion of regeneration, similar numbers of MuSC are present in adult and in aged mice (Fig. 1A).

**Figure 1:**
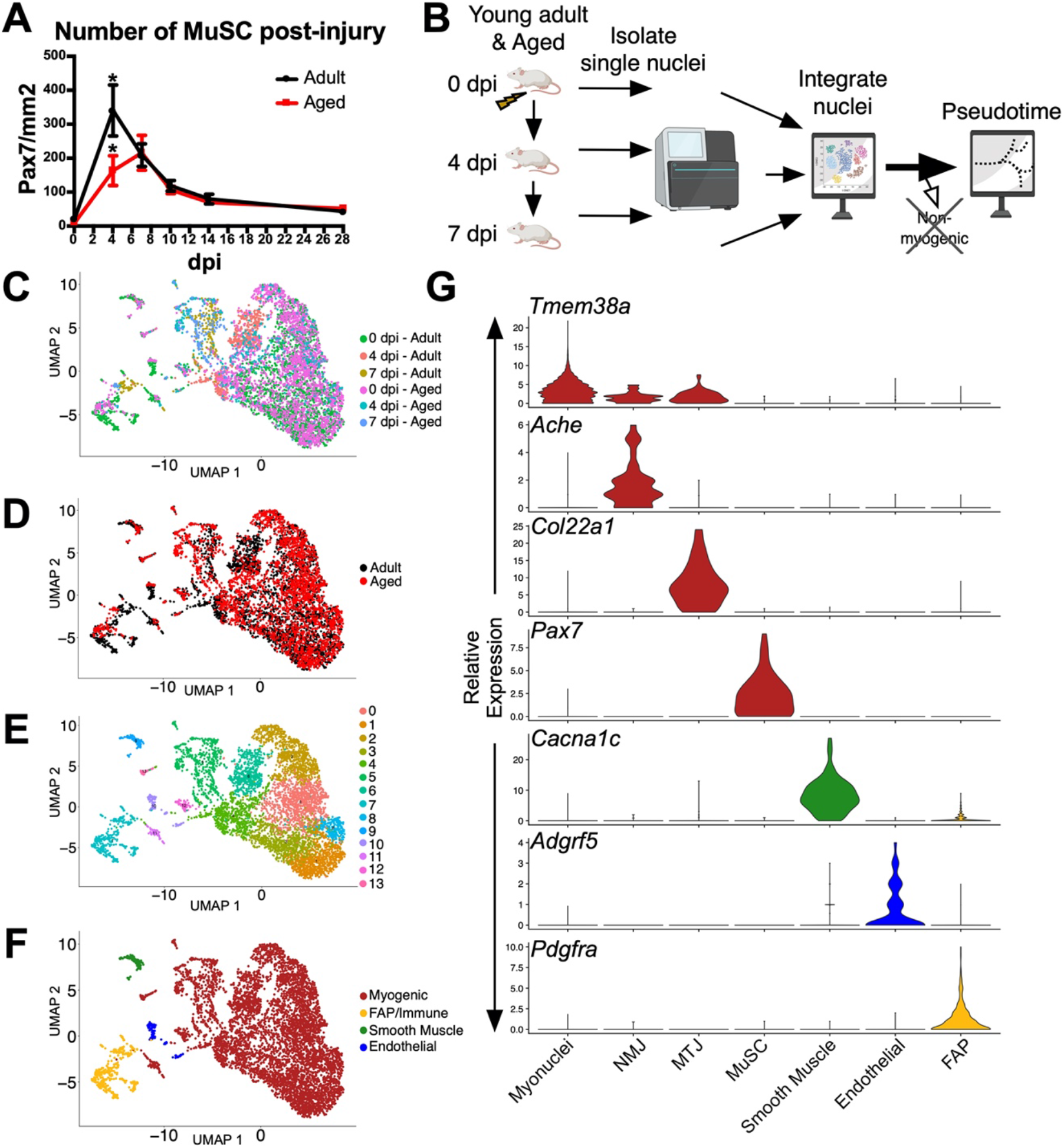
snRNA-seq of regenerating TA muscle from adult and aged mice. (A) Quantification of MuSCs (Pax7+) in the TA muscle during regeneration in young adult (black) and aged (red) mice. \**q* < 0.05. (B) Experimental and computational workflow where single nuclei from TA muscle of adult and aged mice were isolated and enriched for myogenic nuclei at 0 dpi (uninjured), 4 dpi, or 7 dpi, sequenced using the 10X genomics platform, the sequencing data computationally integrated, and clustered by UMAP in *Seurat*. (C) UMAP clusters of nuclei colored by experiment. (D) UMAP clusters of nuclei labeled as either derived from aged (red) or young adult (black) murine TA muscle. (E) *Seurat* initially derived 13 nuclear clusters which were reduced to (F) a single myogenic cluster and 3 others for further analysis. (G) Violin plots of expression levels for individual genes distinguishing nuclear identity.

### snRNA-seq time course in aged and adult mice identifies skeletal muscle nuclei

To explore the transcriptional dynamics occurring during the differentiation of MuSCs into myonuclei and myonuclear maturation, we sequenced single-nuclei (snRNA-seq) from the TA muscle prior to injury, at 4 dpi, and at 7 dpi (Fig. 1B-C). TA muscle nuclei were enriched for myonuclei to increase coverage of low-abundant myogenic populations present during regeneration (27). To assess the coverage of the nuclear isolation and snRNA-seq experiments, the nuclear transcriptomes from each experiment were computationally aggregated to assess the proportions of nuclei derived from mononuclear cells and multinucleated myofibers (8, 28, 29). Nuclei isolated from young adult and aged mice clustered similarly, indicating no major differences between their transcriptional profiles (Fig. 1D). To focus our analysis on myogenic differentiation, 13 *Seurat*-derived clusters were aggregated into a single major myogenic cluster and 3 additional clusters comprised of mononuclear cell nuclei including fibroadipogenic progenitors (FAPs), endothelial cells, and smooth muscle cells (Fig. 1E-G; SI Appendix, Fig. S1A-J) (8, 30). Nuclei comprising the myogenic cluster express either *Pax7* identifying quiescent MuSCs, *Myod1* or *Myog* identifying myogenic progenitors, or *Ckm* or *Mylk2* identifying myonuclei (31–35). The majority of nuclei in the myogenic cluster are myonuclei from myofibers as they express *Tmem38a, Ttn*, *Ckm*, and *Neb* (27, 33, 36) (Fig. 1G; SI Appendix, Fig. S1A-D). MuSCs and myonuclei are discriminated by *Tmem38a* expression, a gene uniquely expressed by myonuclei (Fig. 1G) (27). Among myonuclei, distinct subsets were observed expressing NMJ-associated transcripts and MTJ-associated transcripts, as well as *Myh1, Myh2,* and *Myh4* isoforms, representative of type-IIx, type IIa, or type IIb myofibers, respectively (Fig. 1G; Fig. S1E-J) (6, 8, 11, 17, 37).

### Pseudotime analysis infers a trajectory of progenitor nuclei to mature myonuclei

To characterize the transcriptional changes occurring as MuSCs expand as myoblasts and their differentiation and maturation into myonuclei, we employed *Monocle*, a tool for inferring transcriptional trajectories from single cell sequencing data (38, 39). The trajectory produced by *Monocle* depicts a continuous path for the differentiation of MuSCs into myoblasts, and eventually into mature myonuclei (Fig. 2A). The first of three primary branches in this trajectory consists of MuSCs and myoblasts enriched for *Pax7* and *Ncam1* expression, as well as immature myonuclei expressing *Myh3* and *Myh8*, two immature myosin isoforms (Fig. 2B-F) (16, 31, 40). In the second half of branch 1, and throughout all of branch 2 and branch 3 are differentiated myogenic nuclei expressing *Ttn* and *Neb*, genes common to all myocytes, immature, and mature myonuclei (Fig. 2G-H) (36).

**Figure 2:**
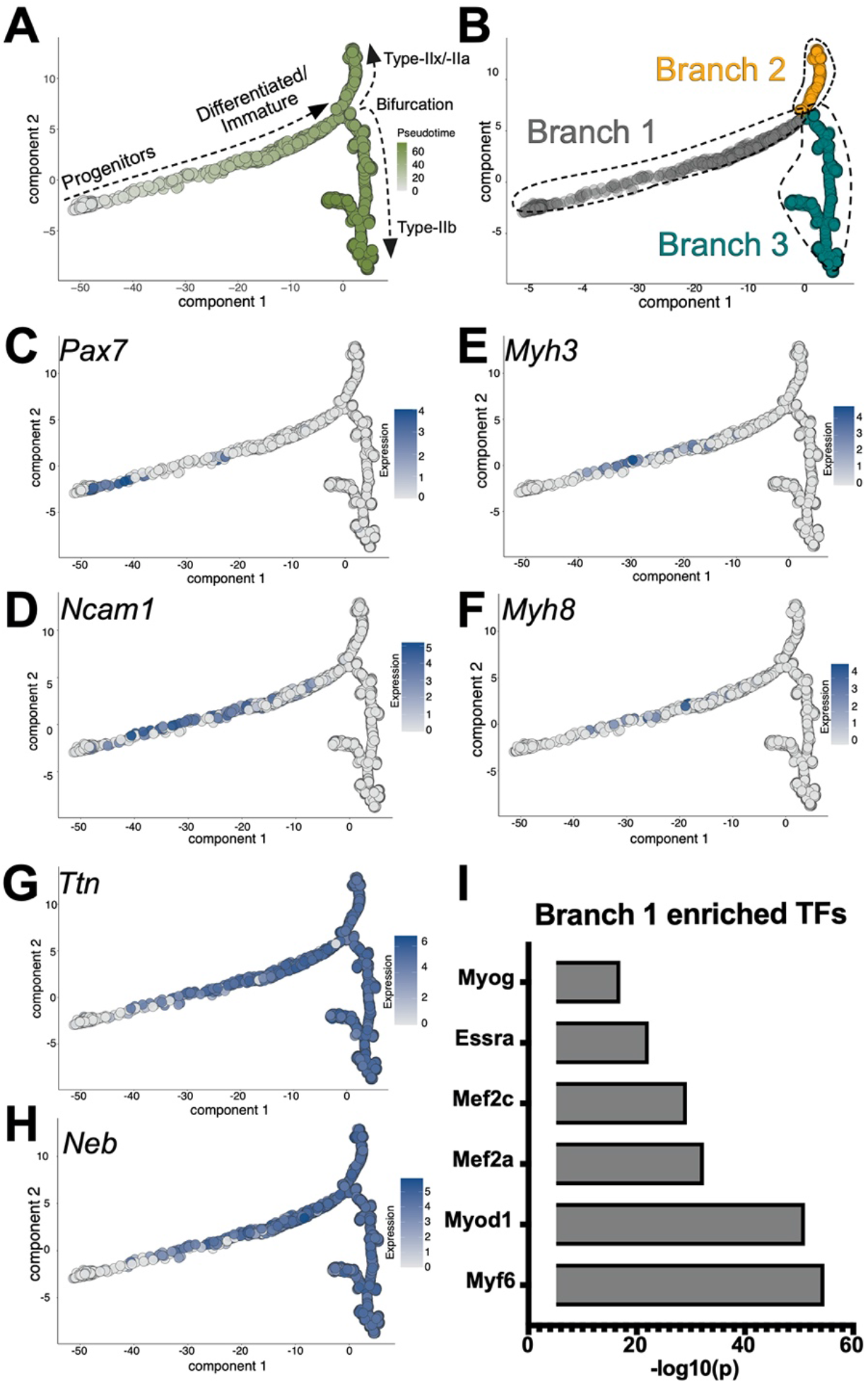
Computationally derived pseudotime trajectories of aggregated myonuclear sequences. (A) *Monocle*-derived pseudotime trajectory of myogenic nuclei during skeletal muscle regeneration (color intensity indicates corresponding pseudotime value). Progenitors on the left differentiate into differentiated/immature myonuclei before bifurcating into the type-II myonuclear subtypes. (B) Pseudotime trajectory with individual branches labeled. (C) Pseudotime trajectories colored in intensity for Expression of myogenic progenitor gene expression *Pax7* and (D) *Ncam1*, (E-F) for immature myonuclei (*Myh3* and *Myh8)*, and (G-H) for universally expressed myonuclear genes (*Ttn* and *Neb*). (I) Myogenic transcription factors enriched for regulating genes specific to branch 1.

MyoD, Myog, Myf6, Mef2a, and Mef2C are myogenic transcription factors regulating gene expression programs maintaining myogenic cell fate and driving myogenic differentiation (32, 41, 42). Whereas MyoD promotes the transition from a quiescent to proliferating myoblasts, Myog expression commits myoblasts to terminal differentiation (32). If branch 1 is comprised of MuSCs, myoblasts, and immature myonuclei then these five myogenic transcription factors should be active in branch 1. We examined the genes enriched in branch 1 for transcription factors that regulate their expression and found that the myogenic transcription factors MyoD, Myog, Myf6, Mef2a, Mef2C, are indeed enriched as regulators of branch 1 gene expression (Fig. 2I). In addition to these five transcription factors is Essra, another myogenic factor that regulates metabolic genes during myocyte differentiation (43).

Gene expression data from the *Monocle*-generated trajectory identifies MuSCs, myoblasts, and immature differentiated nuclei occupying branch 1, while mature and differentiated myogenic nuclei occupy branch 2 and branch 3 (Fig. 2A). We asked if *Monocle* was capable of constructing pseudotime trajectories using only nuclei from injured muscle (data not shown) but found these trajectories failed to distribute nuclei appropriately in pseudotime based on their expected cell fate changes. Therefore, incorporation of nuclei from uninjured and injured muscle into a pseudotime framework is required to generate a trajectory that is consistent with the progression of expected cell fate changes occurring during skeletal muscle regeneration.

### Identification of multiple myogenic progenitors

The presence of a single path of differentiation represented by branch 1 suggests MuSCs, myoblasts, and immature myonuclei all proceed through a uniformed set of transcriptional changes before bifurcating into distinct myonuclear populations. We expected *Pax7* expressing MuSCs to be present at the initiation of branch 1 but instead *Pax7* expressing nuclei were observed just distal to branch 1 initiation (Fig. 2C). The nuclei occupying the proximal region of branch 1 do not express *Pax7* but instead express *Pdgfra*, *Col5a3*, and *Col3a1*, genes expressed in fibroadipogenic progenitor (FAP) cells (8, 44) (Fig. 3A-B). These three FAP-specific genes as well as at least 7 additional genes expressed in this subset of FAP nuclei (SI Appendix, Fig. S2A-G) are expressed in a *Twist2*+ non-myogenic progenitor population that contributes myonuclei to skeletal muscle during maintenance and regeneration (45). This *Twist2*+ population is initially identified by their *Twist2* expression but downregulate *Twist2* before acquiring *Pax7* expression *en route* to differentiating into myonuclei of type-II myofibers (45).

**Figure 3:**
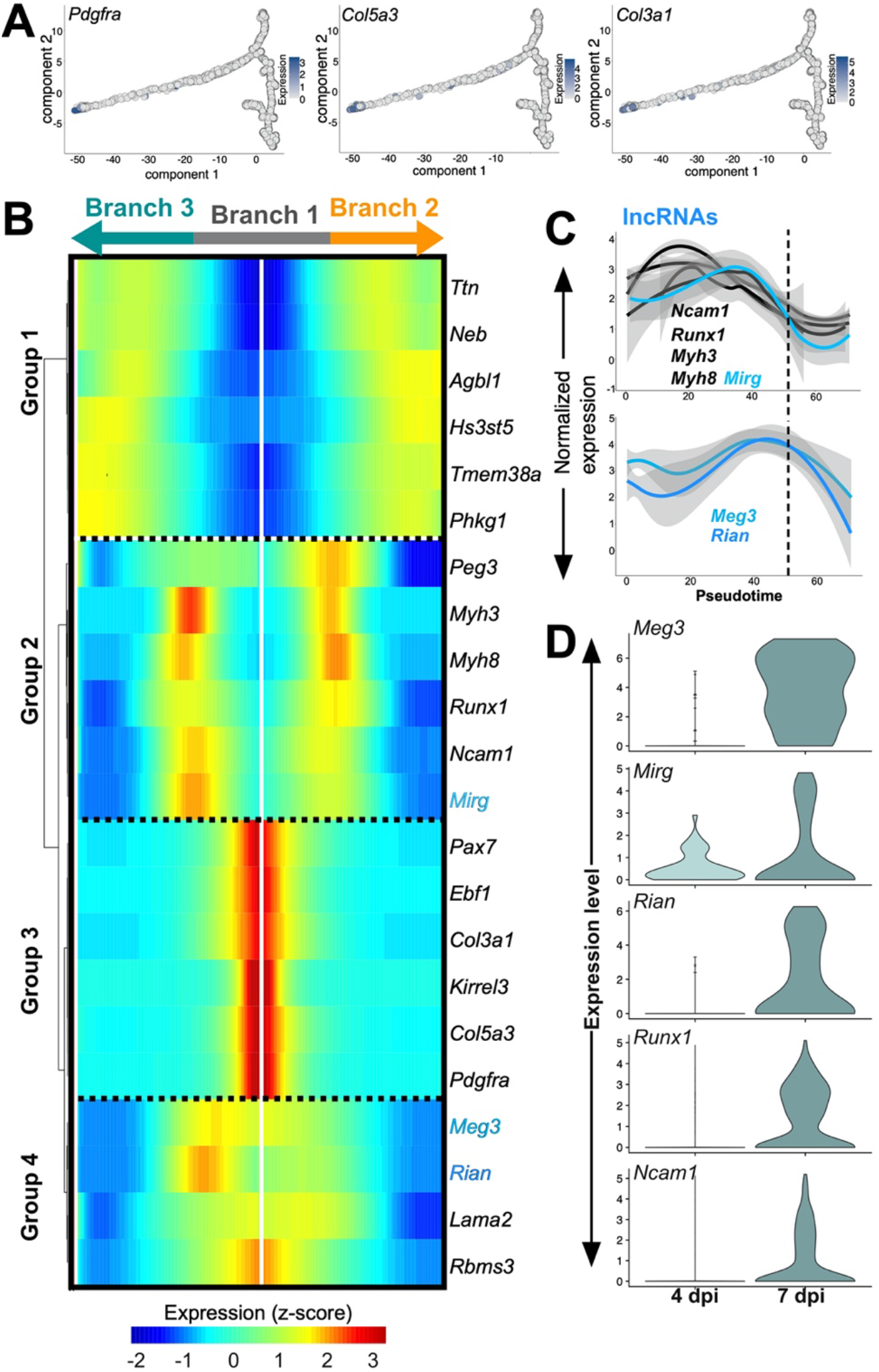
Myogenic progenitor subpopulations. (A) Expression of 3 genes in a non-MuSC FAP/myogenic progenitor population (*Pdgfra, Col5a3, Col3a1)* present in the myogenic pseudotime trajectory. (B) Heatmap depicting the expression of progenitor genes over pseudotime. Pseudotime progresses from center of the heatmap with left and the right representing pseudotemporal expression dynamics in mature myonuclei of branch 2 and branch 3, respectively. Branch markers on top not drawn to scale. Groups are described in text (C) Plots of the pseudotemporal expression distribution for genes expressed in progenitors and the 3 lncRNAs. Dotted line depicts roughly where branch 1 diverges into branch 2 or branch 3. Upper panel, *Ncam1*, *Runx1*, *Myh3*, *Myh8* (black) or *Mirg* (blue). Bottom panel, *Meg3* and *Rian* (D) Violin plots of the relative expression levels at 4 dpi or 7 dpi for the 3 lncRNAs compared to *Ncam1* and *Runx1*.

The pseudotime trajectory depicts the nuclei at the commencement of branch 1 expressing both FAP-associated and myogenic identity genes yet no *Twist2* expression, agreeing with the prior published data where the *Twist2* expressing cells lose *Twist2* as they acquire *Pax7* expression (Fig. 2A; SI Appendix, Fig. S2I). In addition to the 10 genes observed both in the pseudotime trajectory nuclei and the *Twist2*+ population, we identified at least 30 more genes enriched exclusively in the trajectory nuclei. GO analysis of genes enriched in the trajectory nuclei revealed they are indeed a class of fibroblasts (SI Appendix, Fig. S2I; Dataset S1-S2) (46). Thus, the nuclei at the initiation of the pseudotime trajectory are a non-MuSC myogenic progenitor cell population expressing transcripts consistent with FAP and myogenic lineages. Once acquiring myogenic gene expression, the transcriptional trajectory for this FAP subset becomes indistinguishable from the transcriptional trajectory of MuSCs transitioning to myoblasts and differentiating into myonuclei.

### Pseudotemporal characterization of progenitor gene expression

*Monocle* assignments of pseudotime values to individual nuclei enables a deep analysis of gene expression dynamics during regeneration of the TA muscle. All muscle progenitors are present in branch 1, which bifurcates into distinct myonuclear populations in branch 2 and branch 3. The bifurcation of branch 1 occurs after myogenic cells have committed to terminal differentiation and likely have fused into existing myofibers or to generate new myofibers prior to maturing into branch 2 and branch 3 myogenic nuclei.

Genes primarily expressed in branch 1 that differ in expression or transcriptional dynamics in branch 2 compared to branch 3 may expose which genes specify the distinct myonuclear fates in these terminal branches and contribute to the pseudotime bifurcation. Gene expression values spread over a normalized pseudotime scale were plotted as a heatmap to model the dynamics of branch 1-enriched genes from the start of pseudotime to each terminal branch separately (Fig. 3B). Included in this heatmap are 4 groups of genes, where genes in Group 1 exhibit comparable dynamics between branch 2 and branch 3 of the trajectory. Genes in Group 3 are myogenic genes and FAP/myogenic progenitor-specific genes that likewise display similar transcriptional dynamics leading out to branch 2 and branch 3. Given the comparable expression levels and dynamics of genes in genes in Group 1 and Group 3, these two clusters of genes are unlikely to drive the bifurcation occurring at the end of branch 1. In contrast, genes clustered into Group 2 and Group 4 are genes that possess asymmetric expression dynamics with respect to branch 2 or branch 3 transcription. The initial acquisition of myonuclear heterogeneity driven by expression of Group 2 and Group 4 genes within a homogenous pool of progenitors likely propels the divergence of myonuclear populations occupying branch 2 and branch 3.

*Pax7* expression decreases rapidly in branch 1 given its expression is restricted to quiescent and proliferating MuSCs as well as the transitioning FAP/myogenic progenitors, thus clustering into Group 3 with FAP/myogenic progenitor-specific genes (Fig. 3B). Surprisingly, *Myh3* and *Myh8*, generally accepted to be universally expressed as myonuclei mature (16), possess asymmetric expression dynamics with respect to branch 2 and branch 3 and are accordingly clustered into Group 2 (Fig. 3B).

Additionally, three lncRNAs, *Meg3*, *Rian*, and *Mirg*, all transcribed from the common *Dlk1-Dio3* gene cluster (47), are clustered into Group 2 (*Mirg*) and Group 4 (*Meg3, Rian*) (Fig. 3B). All three lncRNAs are involved in muscle development, MuSC differentiation, and are dysregulated during aging and in altered in diseased skeletal muscle (14, 47–49). Although adult myonuclei express *Rian*, there are still conflicting reports regarding whether this locus is active during muscle regeneration (14, 47). LncRNAs are multidimensional regulators of large transcriptional networks (50), thus *Meg3*, *Mirg*, and *Rian* may be key regulators of heterogeneous transcriptional programs during myogenic differentiation.

Extracting the pseudotemporal distributions of *Meg3*, *Rian*, and *Mirg* to visualize their dynamics during TA muscle regeneration better distinguishes their individual profiles (Fig. 3C). The pseudotime profile for *Mirg* in Group 2 is most similar to the expression profiles of *Ncam1* and *Myh3* (Fig. 3C) whereas *Meg3* and *Rian* are distinct from *Mirg*, as *Meg3* and *Rian* both peak in expression closer to the bifurcation point than *Mirg* (Fig. 3C). Regenerating myogenic nuclei at 4 dpi and at 7 dpi express *Mirg,* whereas *Meg3*, *Rian*, *Runx1*, and *Ncam1* are primarily expressed in nuclei at 7 dpi (Fig. 3D). Thus, the genes clustered in Group 2 and Group 4 exhibit asymmetric pseudotime expression in branch 2 and branch 3, suggesting they are part of transcriptional programs that may be drive the acquisition of myonuclear heterogeneity during regeneration.

### The pseudotime trajectory bifurcates into two branches

Branch 1 bifurcates into a shorter branch 2 and a longer branch 3 that distinguish myonuclei comprising type IIa/x or type-IIb myofibers, respectively (Fig. 4A, see Fig. 2A). Nuclei in branch 2 and branch 3 express *Ttn*, *Ckm*, and *Tmem38a* (Dataset S3) and are thus differentiated, yet among the most differentially expressed genes between branch 2 and branch 3 are *Myh1*/*Myh2* and *Myh4,* respectively, revealing the bifurcation point at the terminus of branch 1 segregates myonuclei according to different type-II skeletal muscle subtypes (Fig. 4A; Dataset S4) (10, 13, 19, 46). Although expression of distinct *Myh* isoforms distinguishes branch 2 from branch 3, a cohort of additional transcripts further defines each terminal branch as observed in a pseudotime heatmap clustered into 3 groups (Fig. 4B). Group 1 consists of genes whose expression is symmetric in branch 2 and branch 3, while the Group 2 and Group 3 clusters contain genes whose transcripts are asymmetrically expressed between branch 2 and branch 3, respectively (Fig. 4B).

**Figure 4:**
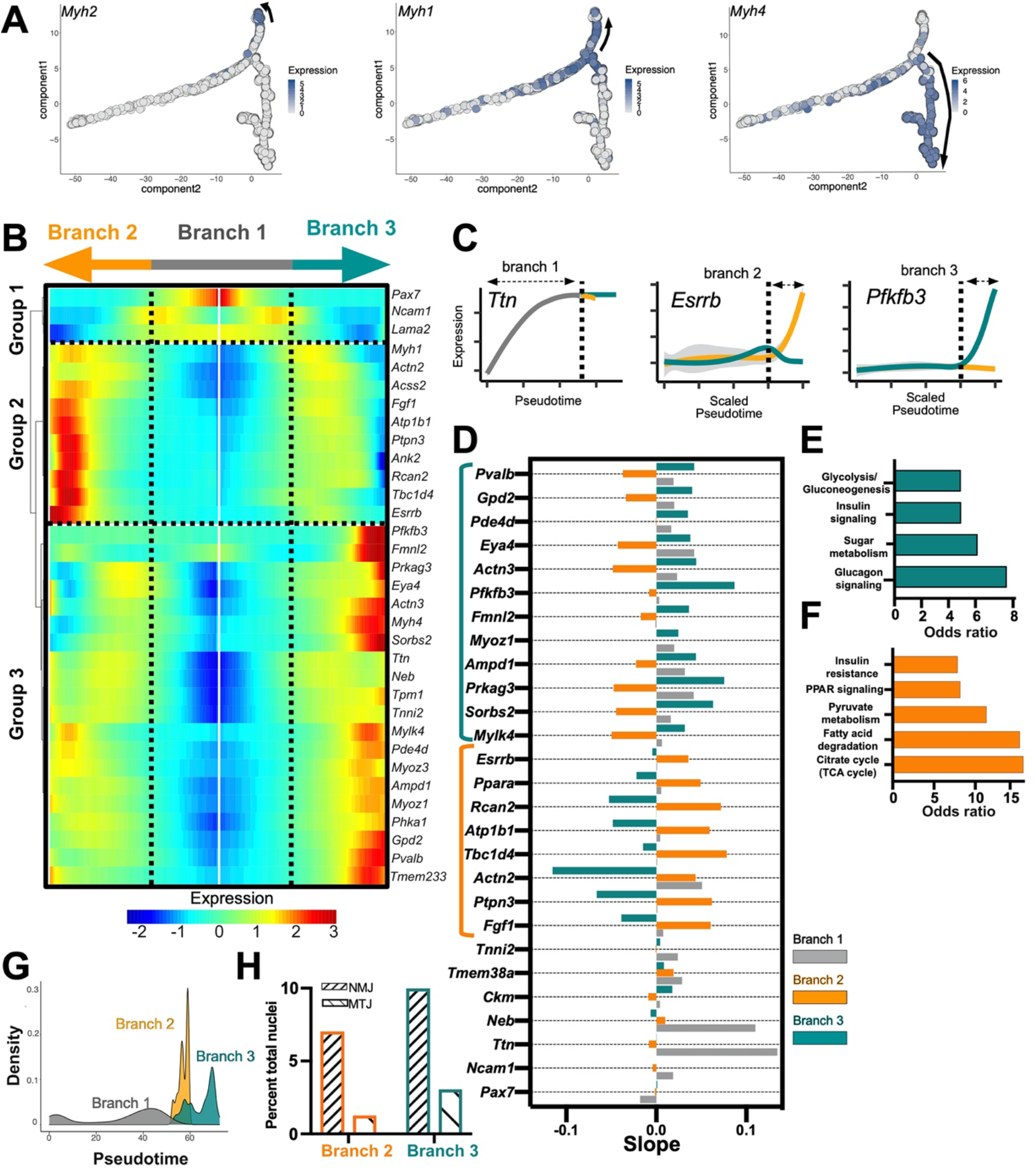
Pseudotemporal analysis of myonuclear gene expression in TA muscles from aged and young mice. (A) Expression of different *Myh* isoforms associated with various contractile speeds. (B) Heatmap depicting pseudotemporal gene expression patterns for mature myonuclei where the center is the start of pseudotime and the left and right represent branch 2 and branch 3, respectively. Progenitor genes are clustered in Group 1, followed by branch 2 and branch 3-specific genes in Group 2 and Group 3, respectively. Genes selected for the heatmap were from differential expression tests between branches and from published snRNA-seq data sets (Petrany et al., 2020). (C) Schematic of how pseudotime slopes were calculated. Genes with positive slopes are increasing expression over pseudotime in a particular branch. (D) Barplot of slopes for individual genes. Left brackets indicate genes that are specific to branch 2 or branch 3. (E-F) Enriched GO categories for genes up-regulated in either branch 2 (lower) or branch 3 (upper). (G) Density histogram of myonuclear psuedotime distributed across the 3 branches. (H) Barplot depicting the proportions of nuclei expressing NMJ- or MTJ-associated genes in branch 2 and branch 3 ((*Ache*, *Chrne*, *Chrna1* (NMJ) or *Col22a1* (MTJ)).

### Pseudotemporal characterization of myonuclear gene expression

To characterize the expression dynamics of genes that define the individual branches, slopes were calculated to quantify the pseudotemporal distributions associated with branch-specific genes revealing three general patterns. The first pattern corresponds to transcripts that are induced in branch 1 but plateau in expression in branch 2 and branch 3. The second are transcripts that are induced primarily in branch 2, while the third represents genes that are primarily upregulated in branch 3 (Fig. 4C), An increase in transcript expression from the start to the end of a particular branch yields a positive slope, while genes neither increasing nor decreasing expression in an individual branch are thus not part of that branches’ transcriptional trajectory and will yield a zero or negative slope (Fig. 4C; SI Appendix, Fig. S3–S4; SI Appendix, Table S1). The pan-myonuclear genes *Ttn* and *Neb,* though are upregulated in branch 1 identified by high positive slopes in this branch, plateau in expression across both branch 2 and branch 3, yielding low or slightly negative slopes (Fig. 4D). In contrast, genes whose transcripts increase in either branch 2 or branch 3 may drive the terminal stages of myonuclear maturation and acquisition of heterogeneity.

We conducted GO analyses of genes enriched in branch 2 versus branch 3 to understand whether the transcripts primarily present in each branch were associated with common biological ontologies (51). The gene expression profiles of myonuclei occupying branch 2 regulate fatty acid degradation and oxidative metabolism (Fig. 4E), while in contrast, branch 3-gene expression profiles are enriched for glycolysis and insulin signaling (Fig. 4E-F; Datasets S6-S7). The segregation of metabolic transcripts for oxidative metabolism in branch 2 and glycolytic metabolism in branch 3 are surprising as both branches are comprised of myonuclei that express myosin isoforms consistent with fast contractile speeds (52), suggesting that these terminal pseudotime branches are comprised of type-II skeletal muscle myonuclei with distinct metabolic states.

The myonuclei in branch 2 and branch 3 appear completely segregated yet myonuclei occupying branch 2 appear earlier in pseudotime than myonuclei in branch 3, with less complexity and fewer additional minor branch points (Fig. 4G). Moreover, NMJ and MTJ myonuclei identified by distinct transcription associated with their anatomical location do not segregate into distinct branches or sub-branches but are similarly represented in branch 2 and branch 3, demonstrating that the anatomically specialized transcripts do not contribute to nuclear segregation in pseudotime (Fig. 4H). Additionally, this supports that branch 2 myonuclei are likely not precursors to branch 3 myonuclei, rather each branch represents transcriptionally and metabolically distinct myonuclear maturation states.

### Pseudotemporal comparisons of myogenic nuclei from adult and aged mice

The pseudotime trajectory was generated by computationally aggregating single nuclear sequencing data from young adult TA muscle and TA muscle nuclei from aged mice (see Fig. 1B). No age-dependent branches were inferred by *Monocle*, demonstrating that the transcriptional trajectories for myonuclear differentiation in young adult and aged mice are similar. However, myogenic nuclei from aged and young adult muscle are differentially distributed across the individual trajectory branches (Fig. 5A). Fewer myogenic nuclei from aged mice occupy branch 1 compared to adult mice while three quarters of the nuclei in branch 2 are from aged mice, and roughly equal proportions are present in branch 3 (Fig. 5B). The differential pseudotime distributions for myogenic nuclear transcripts occurring between aged and young adult muscle does not arise from a lack of sampling as equal numbers of nuclei were assessed for gene expression (Fig. 5B). Thus, the pseudotime distribution of transcripts expressed in myogenic nuclei from regenerating TA muscles of aged mice are distinct from those in young adult mice, producing three quarters of the branch 2 trajectory (Fig. 5B).

**Figure 5:**
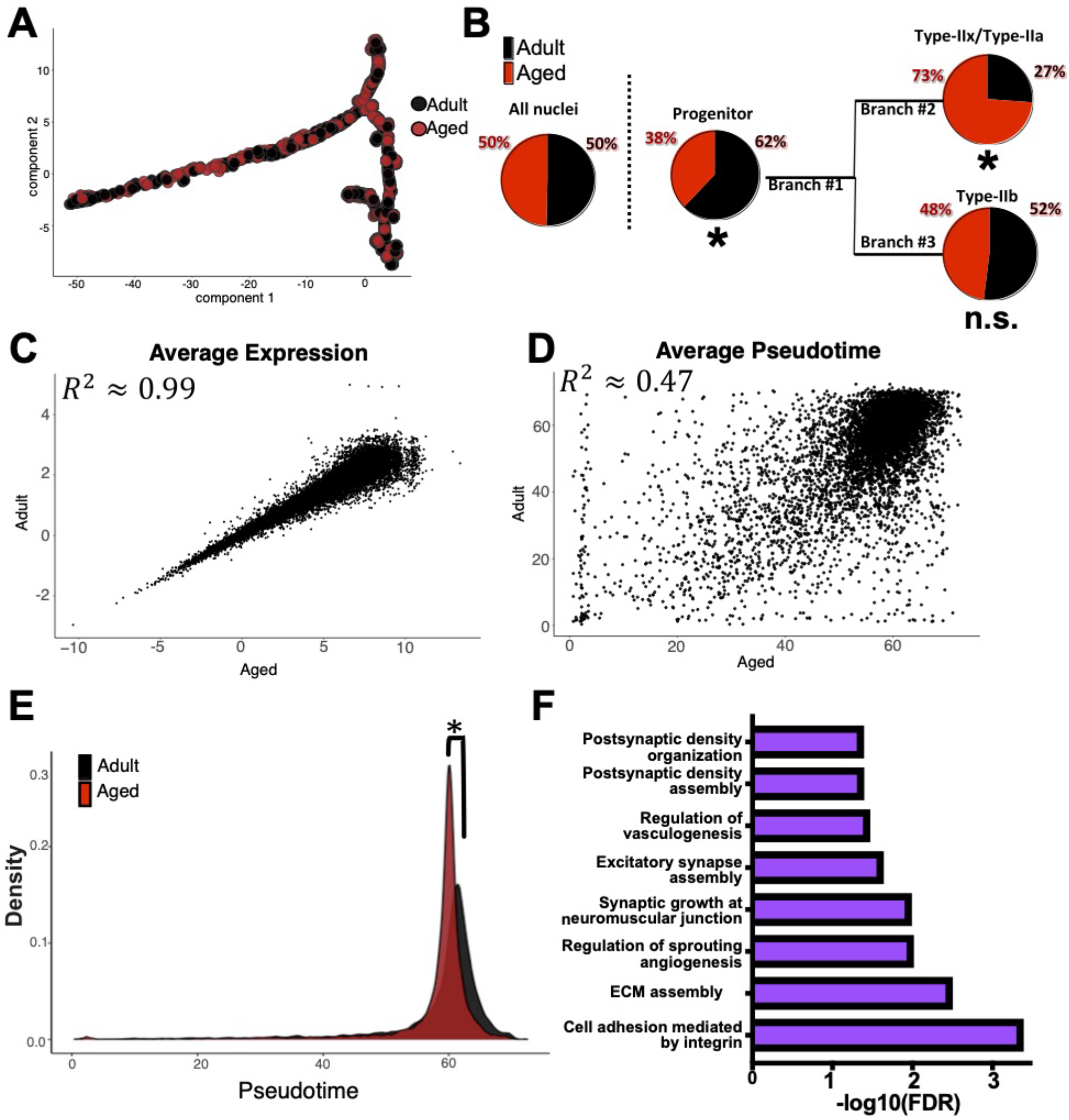
Pseudotemporal comparisons of myogenic nuclei from adult and aged mice reveals similarities and differences. (A) Pseudotime trajectory of nuclei from aged and adult regenerating skeletal muscle. (B) Pie charts depicting the proportions of nuclei from adult or aged muscle distributed across all 3 branches. \**p* < 0.05, Chi-squared test. (C) Averaged normalized expression, (D) or pseudotime, values for individual genes (points) from either aged or adult muscle nuclei. (E) Density histogram of pseudotime distributions for nuclei from aged and adult muscle, \**p* < 0.05 Wilcoxon rank-sum test. (F) Enriched GO categories for the top 1000 genes with largest pseudotemporal shifts in myonuclei of TA muscles from aged mice and adult mice.

The aggregated data for nuclear transcriptomes from young adult and aged TA muscles failed to identify age-specific pseudotime branches, suggesting the overall transcriptional levels are similar despite the clear difference in proliferation following an induced injury (See Fig. 1A). Plotting averaged expression levels of mutually expressed genes in young adult myogenic nuclei compared to myogenic nuclei in TA muscles of aged mice the expression levels are very similar, revealing few differentially expressed genes (Fig. 5C). In striking contrast, averaged pseudotime values of gene expression in myogenic nuclei from young adult mice plotted against gene expression in myogenic nuclei from TA muscles of aged mice are highly divergent (Fig. 5D).

During muscle regeneration, MuSC activation and myogenic progenitor expansion is delayed in aged mice compared to adult mice (Fig. 1C). Plotting nuclear pseudotime values as a histogram reveals a delay in pseudotime for gene expression in nuclei from aged TA muscle compared to pseudotime values for gene expression in myogenic nuclei from TA muscles of young mice (Fig. 5E). GO term enrichment for genes in myogenic nuclei from TA muscle of aged mice reveals that genes involved in innervation, synaptogenesis, and vasculogenesis were among the most enriched GO categories pseudotemporally delayed in myogenic nuclei from aged mice compared to adult mice (Fig. 5F; Datasets S8-S9). This is an intriguing observation given the established functional reduction of NMJs in aged muscle (21).

## Discussion

Skeletal muscle myonuclei arise from fusion of myogenic progenitors during development or from skeletal muscle stem cells during regeneration, yet are transcriptionally heterogenous despite sharing a common cytoplasm (7, 8, 12, 13). Following a muscle injury, homogeneous MuSCs activate, proliferate as myoblasts, and then differentiate and fuse to repair and replace damaged myofibers (53, 54). We used *Monocle* to construct a pseudotime trajectory for myogenic nuclear gene transcription during skeletal muscle regeneration, which revealed unexpected contributions of non-myogenic precursors to skeletal muscle and identified the onset of myonuclear transcriptional heterogeneity. A non-myogenic progenitor involved in muscle regeneration identified by *Monocle* matures into myonuclei with transcriptional dynamics indistinguishable from MuSC-derived myoblasts. Second, transcriptionally homogenous myogenic progenitors initiate differentiation and then diverge as myonuclei mature acquiring distinct transcriptional profiles associated with oxidative metabolism and glycolytic metabolism. Finally, myogenic nuclei from aged mice differentially segregates throughout the trajectory, with myonuclei of aged mice preferentially populating the more oxidative branch 2 as opposed to the glycolytic branch 3.

MuSCs are essential for skeletal muscle regeneration producing myonuclei (25), yet the initiation of the pseudotime trajectory is populated by nuclei expressing genes consistent with both FAP cells and myogenic cells. FAPs are a muscle resident mononuclear cell involved in skeletal muscle maintenance and regeneration. FAPs in skeletal muscle are functionally diverse as they are capable of transdifferentiating into myonuclei during muscle development, producing MTJ myonuclei and myonuclei in Type-II myofibers (30, 45, 55, 56). The *Monocle*-inferred trajectory identifies a subset of nuclei expressing FAP and myogenic identity genes just prior in pseudotime to the entry of MuSCs into the trajectory, suggesting that the subset of FAPs observed in the trajectory transitions through a transcriptional state similar to *Pax7*+ MuSCs. Upon this transition, these FAP/myogenic progenitors merge into the MuSC trajectory and are transcriptionally indistinguishable from proliferating myoblasts. Thus, myonuclei are derived from at least two distinct cell types during regeneration but the immature myogenic cells are indistinguishable from each other and comprise a common precursor for all mature myonuclei in the TA muscle.

Mature myonuclei segregate into two major pseudotime trajectories represented by branch 2 and branch 3 that are distinguished by divergent transcriptional profiles including expression of distinct *Myh* isoforms and the expression of a cohort of metabolic genes. The myosin isoforms and metabolic genes expressed by the myonuclei in these distinct branches specify type-IIx/-IIa fast-oxidative myofibers, and type-IIb fast-glycolytic myofibers (7, 8, 13, 52, 57, 58). Surprisingly, myonuclei within branch 2 and branch 3 possess similar gene expression profiles for NMJ and MTJ nuclei, suggesting the specification of NMJ and MTJ myonuclei occurs after the bifurcation of branch 1 into branch 2 and branch 3. Although myofibers are capable of switching contractile types (17, 59, 60), the mechanisms involved are unclear and whether myonuclear accretion is required for transitioning contractile types is unknown. Once nuclear transcriptional profiles bifurcate into branch 2 or branch 3, the transcriptional trajectories terminate in each branch, suggesting interconversion between myonuclei in branch 2 and branch 3 is unlikely. Thus, accretion of new myonuclei from the common myogenic precursor population may be necessary for myofibers interconvert their muscle fiber type.

Skeletal muscle maintenance and regeneration in aged mice as well as in aged humans is compromised in part due to deficits in MuSC function (19, 61). MuSCs from aged mice activate and proliferate with delayed kinetics compared to MuSCs from young mice (62) and in culture, MuSCs from aged mice differentiate rather than continue to proliferate (19). We confirmed these observations *in vivo* with EdU lineage tracing, demonstrating that MuSCs in aged mice are impaired, failing to proliferate and generate the numbers of myoblasts generated in young adult mice following an induced muscle injury. While the overall pseudotime trajectory and gene expression profiles were broadly consistent between myogenic nuclei from TA muscles of aged mice compared to young adult mice, myogenic nuclei from aged mice differentially populated the pseudotime trajectories with branch 2 type-IIa/x myonuclei disproportionately populated with nuclei from aged mice, and myogenic nuclei in branch 1 comprised of a greater proportion from young adult mice. The delay in activation and failure of MuSC progeny to fully expand when TA muscles are injured in aged mice correlates with the proportion of myogenic nuclei from aged mice that preferentially populate branch 2, which is earlier in pseudotime than branch 3. The genes that are most significantly pseudotemporally shifted in myogenic nuclei from aged mice are predominately involved in synapse and NMJ formation and maintenance, providing a potential mechanism underlying compromised innervation in aged muscle. Altogether, alterations in transcriptional dynamics rather than relative gene expression levels may primarily drive degradation of muscle function during aging.

## Materials and Methods

### Mice

Mice were bred and housed according to National Institutes of Health (NIH) guidelines for the ethical treatment of animals in a pathogen-free facility at the University of Colorado at Boulder. University of Colorado Institutional Animal Care and Use Committee (IACUC) approved animal protocols and procedures. Young adult mice were C57B6 mice, (Jackson Labs Stock No. 000664) between 4 and 8 months old and were a mix of male and female. The aged mice used were F1 mice from a C57BL/6J and DBA/2J cross (Jackson Labs No. 100006), collected between 24-28 months old and a mix of male and female. For injuries, mice were anesthetized with 3% isofluorane followed by injection with 50μL of 1.2% BaCl2 into the left TA and EDL muscle.

### Immunofluorescence staining of tissue section

The TA muscle was dissected, fixed for 2h on ice cold 4% paraformaldehyde, and then transferred to PBS with 30% sucrose at 4°C overnight. Muscle was mounted in O.C.T. (Tissue-Tek®) and cryo-sectioning was performed on a Leica cryostat to generate 8 μm(?) sections. Tissues and sections were stored at −80°C until staining. Tissue sections were post-fixed in 4% paraformaldehyde for 10 minutes at room temperature and washed three times for 5 min in PBS. To detect Pax7 on muscle sections we employed heat-induced antigen retrieval. Post-fixed sections on slides were placed in a citrate buffer (100mM Sodium citrate containing 0.05% Tween20 at pH 6.0) and subjected to 6 min of high pressure-cooking in a Cuisinart model CPC-600 pressure cooker. For immunostaining, tissue sections were permeabilized with 0.5% Triton- X100 (Sigma) in PBS containing 3% bovine serum albumin (Sigma) for 30 min at room temperature. EdU was visualized following manufacturers guidelines (ThermoFisher Scientific cat#C10337). Primary antibodies included anti-Pax7 (DSHB) at 2 μg/mL and rabbit anti-laminin (Sigma-Aldrich cat#L9393) at 2.5 μg/mL incubated on muscle sections for 1h at room temperature. Alexa secondary antibodies, donkey anti mouse Alexa Flour 647, donkey anti-rabbit Alexa Flour 555 (ThermoFisher) were used at a 1:750 dilution and incubated with muscle sections for 1 h at room temperature. Prior to mounting, muscle sections were incubated with 1 μg/mL DAPI for 10 min at room temperature then mounted in Mowiol supplemented with DABCO (Sigma-Aldrich) as an anti-fade agent.

### Microscopy and image analysis

All images were captured on a Nikon inverted spinning disk confocal microscope.

Objectives used on the Nikon were: 10x/0.45NA Plan Apo and 20x/0.75NA Plan Apo Images were processed using Fiji ImageJ. Confocal stacks were projected as maximum intensity images for each channel and merged into a single image. Brightness and contrast were adjusted for the entire image as necessary.

### Statistical analysis of Pax7 quantification

Statistical tests were performed on Pax7 quantified cells per mm^2^ for N = 3 young adult and N = 3 aged mice in Prism (GraphPad). Significance was assessed using two-tailed unpaired Student’s t test with two-stage step-up method of Benjamini, Krieger, and Yekutieli multiple comparison adjustment with a *q* < 0.05 considered significant. Each N was generated from an individual mouse.

### Nuclear isolation for sequencing

Nuclei were isolated from single tibialis anterior (TA) muscles with the following modifications (Cutler et al., 2017). The tibialis anterior (TA) muscle was dissected, weighed, and placed in a microcentrifuge tube. The muscle was chopped with scissors (FST Item No.15024-10 in homogenization buffer (10mM HEPES, 60mM KCl, 2mM EDTA, 0.5mM EGTA, 300mM sucrose). The muscle was homogenized (homogenizer info) for 45 strokes and the homogenate filtered using 100uM filters (filter info). Cell strainers were pre-wet with Homogenization Buffer before samples were placed on them. Myonuclei were enriched by differential centrifugation (Ti- 14) for 3hrs at 33,000rpm, and the purified nuclei were resuspended in PBS and 0.1% BSA. 300 Units of RNase OUT was added to prevent RNA degradation. The nuclei counted with a TC20 Automated Cell Counter (Catalog #145-0101). and processed for single nuclear sequencing.

### snRNA sequencing

A total of 6 samples were prepared for snRNA-sequencing: adult uninjured TA (2x), aged uninjured TA (2x), adult 4 dpi (1x), adult 7 dpi (1x), aged 4 dpi (1x), aged 7 dpi (1x). snRNA sequencing was carried out using the 10X Genomics Chromium Single Cell 3’ Protocol, Accessories and Kits (CG000183 Rev A). 1,600 nuclei were loaded per reaction in the chromium controller for a targeted recovery of 1,000 nuclei. The samples were processed following the manufacturers guidelines exactly and sequenced using the Illumina Nextseq500.

### Initial data processing with *Cellranger*

Initial quality control of sequencing was conducted using FastQC (v0.11.8). The CellRanger Software Suite (10x Genomics, v3.0.1) was then used to process single-nuclear FastQ files. For transcriptome alignment, a custom “pre-mRNA” mm10 reference package was used as previously described (8, 63).

### Initial data processing in *R*

All code is publicly available, but briefly, background noise, low quality nuclei, or nuclei derived from doublets, were removed for downstream analyses using *SoupX* 1.4.5 and *Seurat* 3.1.4. Generally, the standard pre-processing workflow for integrating multiple datasets suggested by *Seurat* was used with minor changes to parameters.

Upon dimensional reduction performed by UMAP, cell type identification was carried out using FindMarkers to reveal differentially expressed genes from individual clusters. Genes were cross-referenced with literature, as well as compared to the Myoatlas database webtool for skeletal muscle nuclear gene expression (8).

### *Monocle*/*Seurat*

Before processing *Seurat* objects in *Monocle* 2.14.0, nuclei from each experiment were filtered for their expression of either *Pax7*, *Myod1*, *Myog*, *Ckm*, or *Mylk2* to ensure an entirely myogenic nuclear population. Upon merging myogenic nuclei from each experiment, *Monocle* inferred the pseudotime trajectory containing 28 branch points and 59 states. States were then aggregated based physical location in the trajectory to reduce the branches down to 3. Differential gene tests between branches were run BEAM in *Monocle*, or alternatively data was exported back into *Seurat* and FindMarkers was used to compare nuclei using “state” and “pseudotime” assignments inferred by *Monocle.*

For pseudotime heatmaps the plot_genes_branched_heatmap function in *Monocle* was used, using the Monocle-assigned branch points surrounding the bifurcation point.

### Young vs aged statistics

Pie chart significance was calculated using Chi-square tests with 50% distribution of adult and aged nuclei as the expected values and a df = 5 and \**p* < 0.05. Differences between pseudotime distributions of all myogenic nuclei was assessed using the Wilcoxon rank-sum test in *R*, statistical significance was assessed as having *p* < 0.05.

### Slope analysis

*R* code for calculating slopes is provided, but in short to calculate slopes associated with individual genes within individual branches, nuclei were first plotted by their expression and their *Monocle*-assigned pseudotime values. For branch 1, slopes represent the slope of a line spanning from the start of branch 1 to the bifurcation point at roughly 50 on the original pseudotime scale. For branch 2 and branch 3, we first concatenated lines from branch 1 onto either of these terminal branches, creating a single curve associated with either branch 2 or branch 3 expression of a particular gene. The nuclei in a small sub-branch off branch 3 were removed for this slope analysis. Given that branch 2 was shorter and appeared earlier in pseudotime than branch 3, we scaled distributions for either branch 2 or branch 3 lines to an arbitrary scale of 0-100. From here, slopes were calculated for nuclei after 75 scaled pseudotime units.

### GO analyses

GO analysis was conducted with Panther or Enrichr (64–66). TF enrichment in branch 1 was from the ARCHS4 TF Coexpressed database. Pathways enriched for genes specific to branch 2 compared to branch 3, and vice versa, were from the Kegg 2019 pathways database (51). Enriched ontologies for top 1000 genes pseudotemporally shifted genes were assessed using Panther “GO biological pathway complete 2021” compared to a background gene list of all genes detected in myogenic nuclei (Dataset S9).

## Data and materials availability

All data is available in the main text or the supplementary materials. Code will be available on the Github repository https://github.com/jeku7901-CU/myonuclear_maturation.git and sequencing data deposited in GEO (awaiting accession number).

---

**Figure S1:**
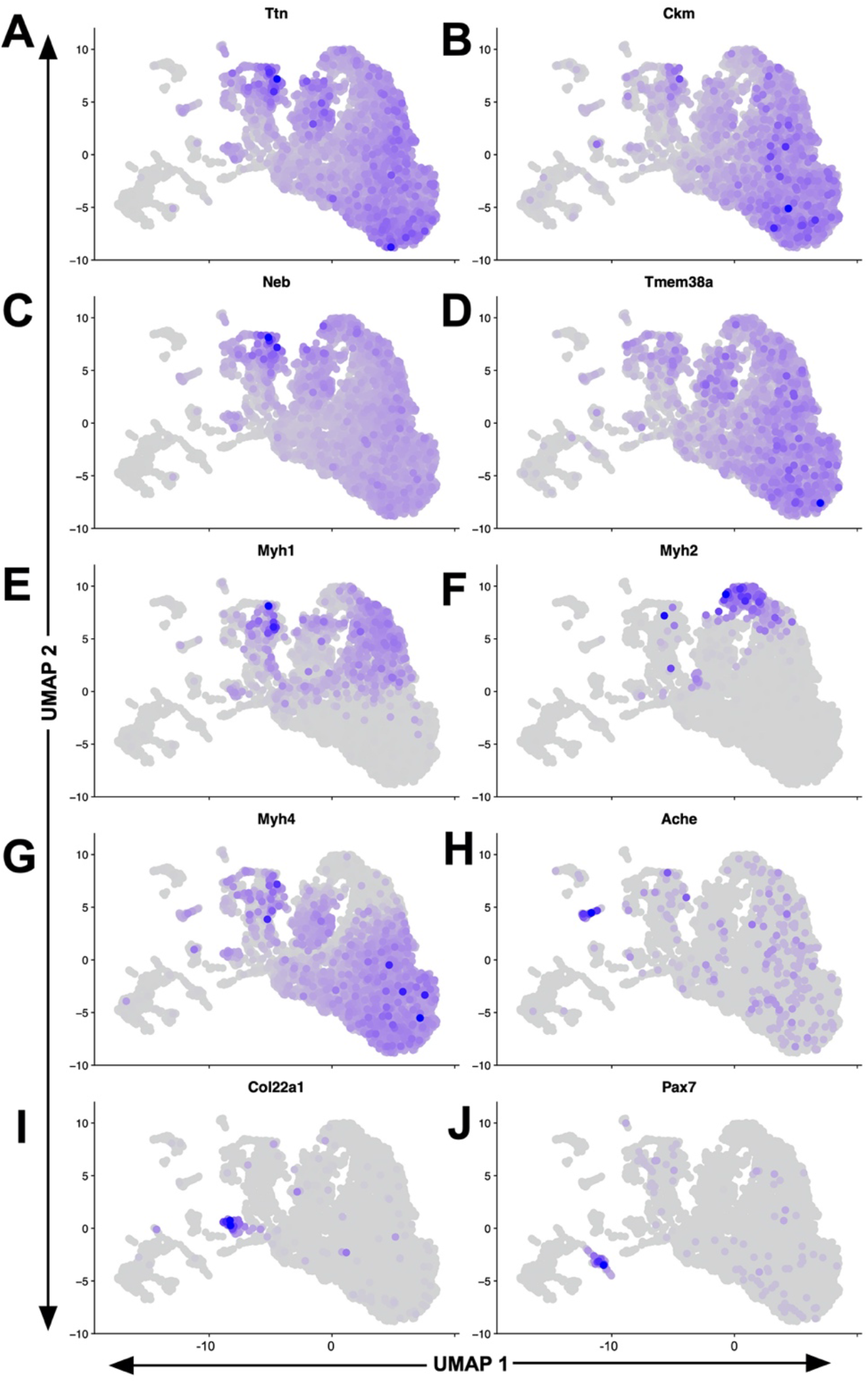
UMAP plots of nuclei colored by normalized expression. (A-J) UMAP plots depicting the expression of individual genes across all nuclei used for nuclear type classifications. Intensity of purple indicates higher expression of genes distinguishing nuclei (corresponds to Figure 1G).

**Figure S2:**
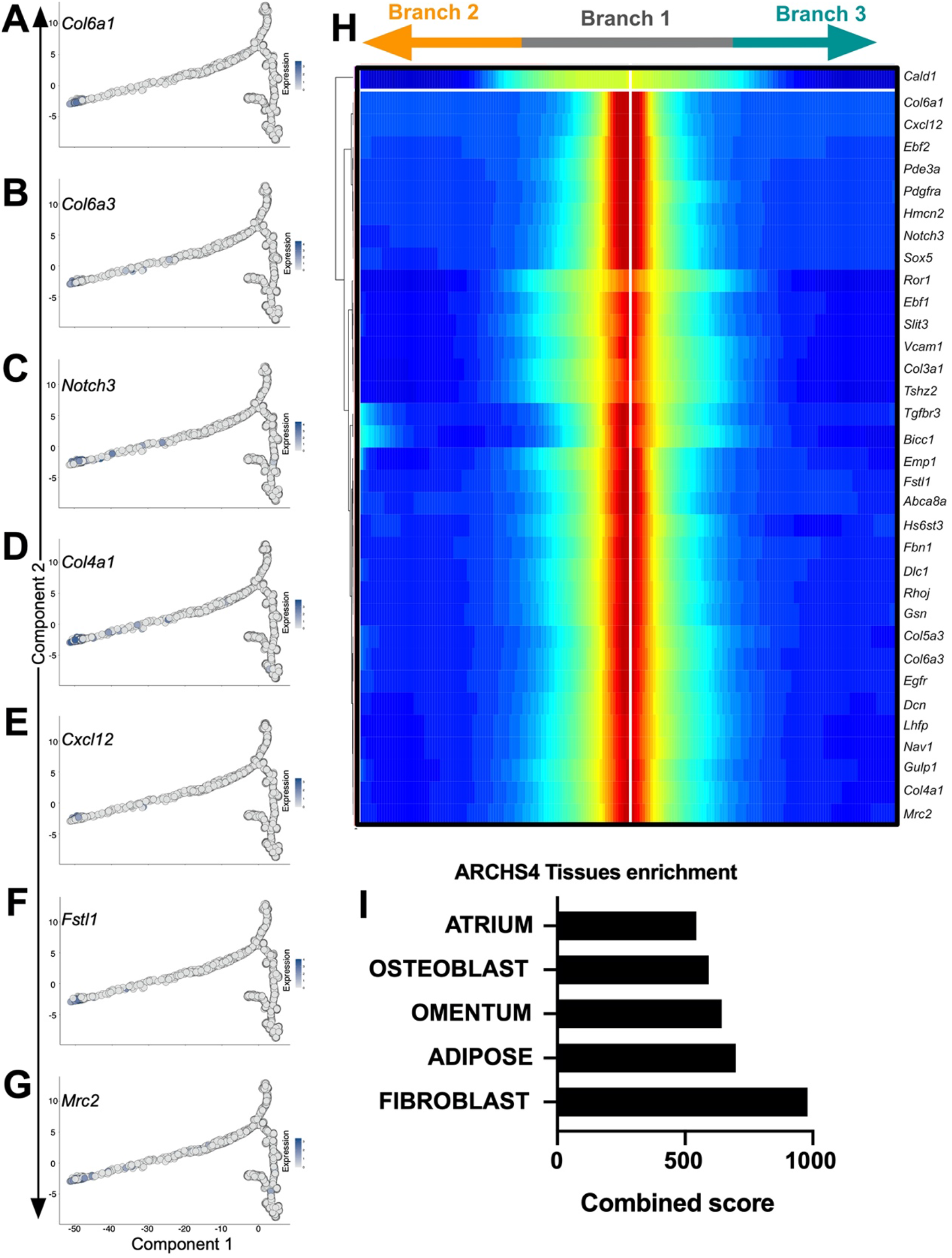
Gene expression associated with mixed FAP and myogenic nuclei. (A-G) Trajectory plots of specific genes expressed in this group of nuclei prior to *Pax7* expressing nuclei. (H) Pseudotime heatmap similar to Figure 3 depicting the expression of genes enriched in this group. Branch notation on top of heatmap is not drawn to scale. (I) GO analysis of genes enriched in this group with cell type enrichment.

**Figure S3:**
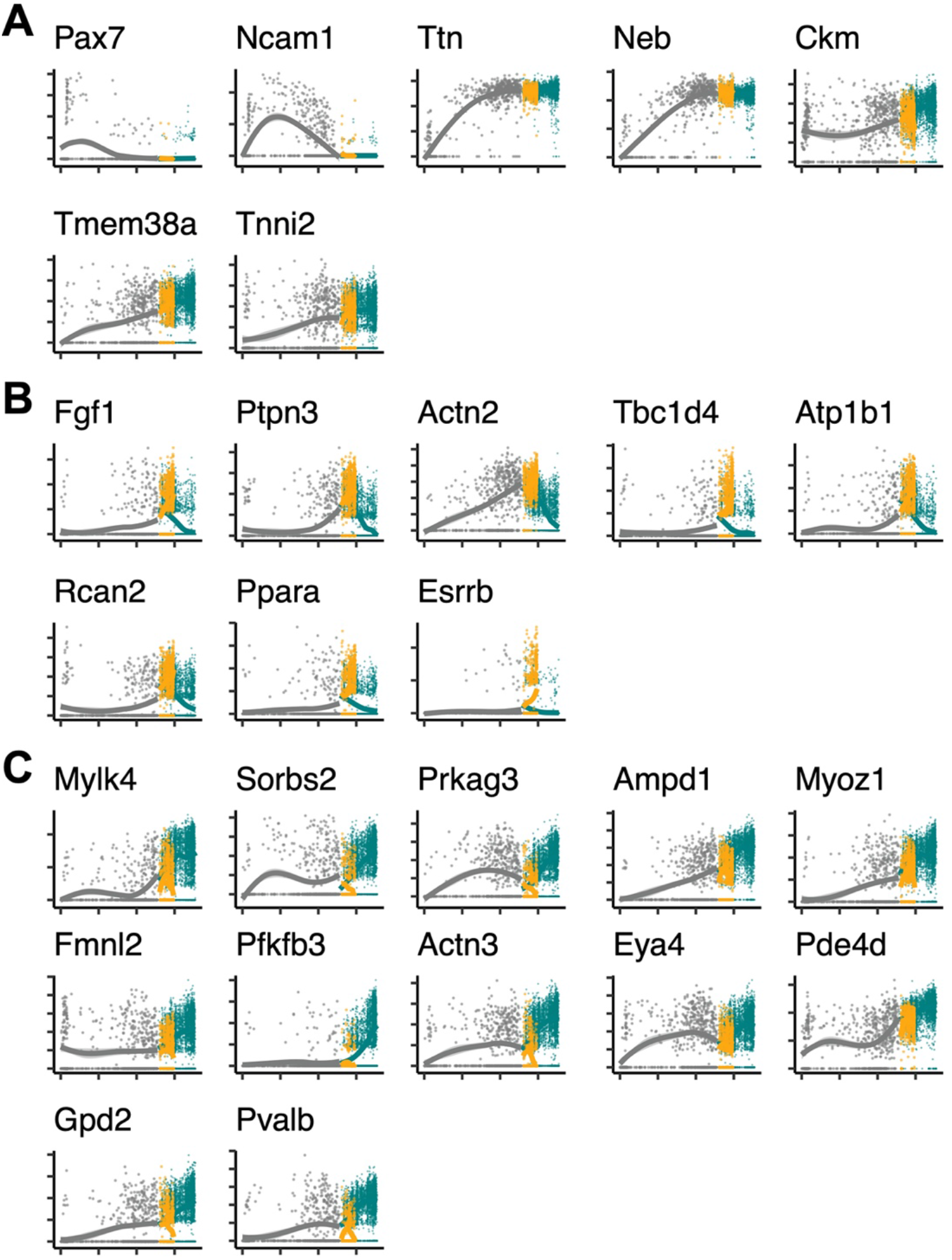
Raw pseudotemporal expression distributions for individual genes. Individual nuclei were plotted for by their expression (y-axis) of the indicated gene and their assigned pseudotime values (x-axis). Nuclei are colored by their corresponding branch from the trajectory. In Figure 4D, branch 1 slopes were calculated as the slope of the nuclei colored gray.

**Figure S4:**
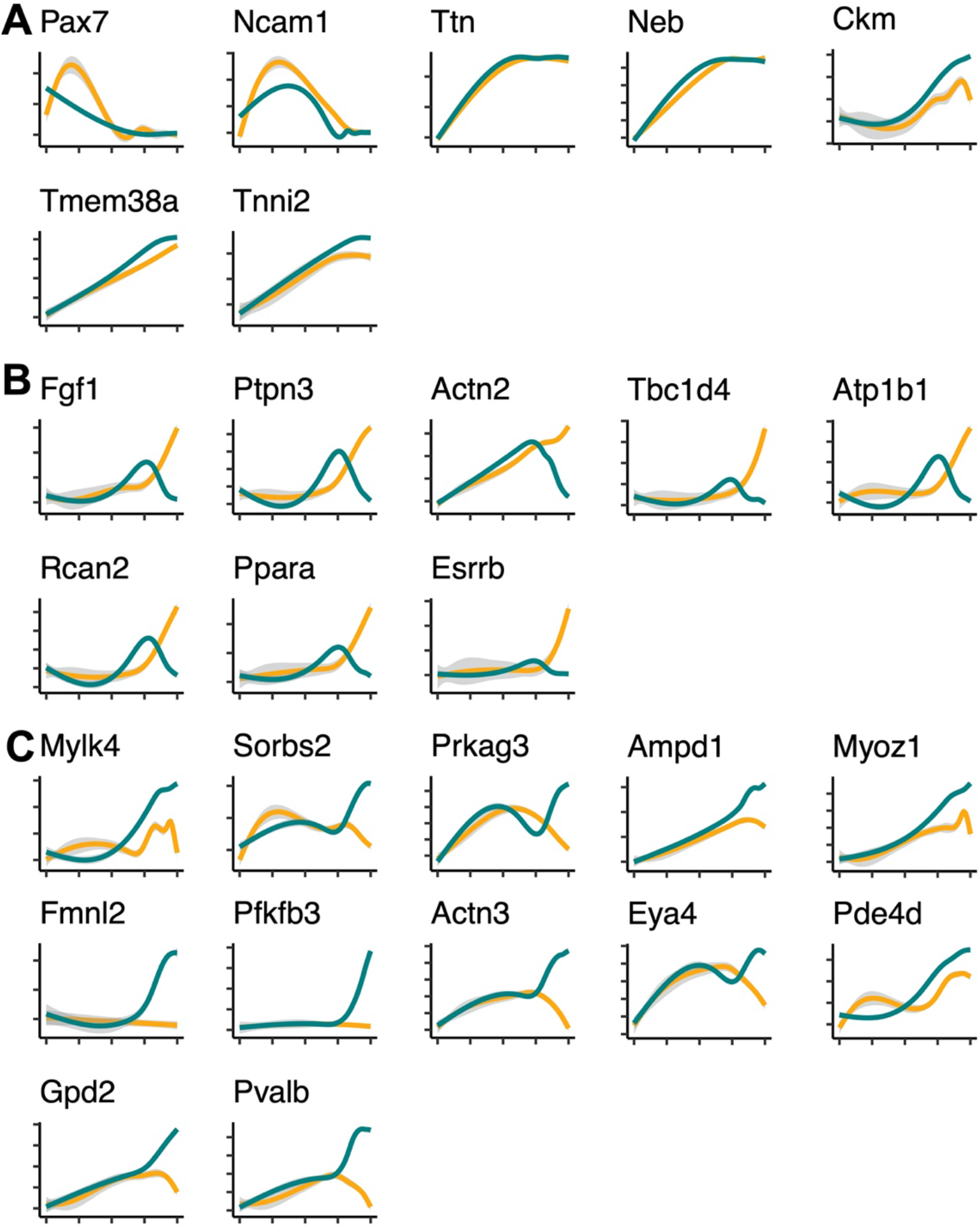
Scaled pseudotemporal expression distributions for individual genes. Expression levels for individual genes (y-axis) plotted as a function of pseudotime (x-axis). Each genes’ pseudotime distribution was linearly transformed to adjust pseudotime range to scaled pseudotime units of 0-100. In Figure 4D, branch 2 and branch 3 slopes were calculated for each line after 75 scaled pseudotime units.

